# SIMHYB 2: a software tool to explore and illustrate evolutionary forces in Population Genetics teaching and research. Application to Conservation Genetics

**DOI:** 10.1101/2024.02.13.580073

**Authors:** Álvaro Soto, David Rodríguez-Martínez, Unai López de Heredia

## Abstract

Practical approaches have become a standard in many scientific disciplines, including population genetics. By analyzing properly selected datasets, the students can calculate parameters and draw conclusions about genetic diversity, differentiation and evolution of populations with higher efficiency than if based exclusively on theoretical lessons. However, preparing the appropriate datasets is a hard task and a wrong selection can spoil a well-aimed practice. Here we present SimHyb 2, a software tool specifically intended to ease the full understanding of evolutionary forces by the students and to help the teacher to prepare adequate datasets and examples for the practices. It simulates the course of a mixed population under user-defined reproductive and evolutionary conditions. Outputs can be easily adapted for downstream analysis with other popular tools as GenAlEx or Structure. Thus, SimHyb 2 is very suitable for project-based-learning approaches: students can produce their own datasets in different scenarios of genetic drift, migration, selective advantage, reproductive success… Additionally, SimHyb 2 is the only simulation software available to date providing traceable pedigrees of individuals, being therefore very convenient for preparing datasets for parentage analysis, spatial genetic structure or conservation genetics study cases. Satisfactory results from its ongoing utilization in higher education and research are reported.

## INTRODUCTION

Genetics and evolution are foundational pillars of life sciences. Population genetics involves the examination and modelling of changes in the frequencies of genes and alleles in populations over space and time and is a topical matter in biology, biotechnology, agronomy and forestry undergraduate and graduate courses. However, the intricacies of genetic concepts and mathematical models can pose challenges for students (Knippels, 2002), who must struggle with probability and statistics underpinning many population genetic models. As in other scientific subjects, the presentation of population genetics concepts only through lectures can give many students just a superficial understanding that may lead to misconceptions (Kontra et al., 2015). Genetic phenomena transverse multiple levels of organization, and reasoning across these levels is challenging for students (van Mil et al. 2013). In addition, many population genetics concepts and ideas are unfamiliar to high school and university students (Duncan & Reiser, 2007), and are sustained on a relatively complicated mathematical-theoretical basis that is not easy to learn through traditional master lessons. More specifically, previous literature has identified several challenging population genetics topics, such as the dynamic nature of genetics (Haskel-Ittah et al., 2020) the molecular basis of genetic variation (Lewis & Katman, 2004), genetic drift (Andrews et al., 2012) or proper comprehension of statistical genetics (Neyhart & Watkins, 2020), among others.

Active learning strategies through case-study resolution and student-driven discussion are pivotal teaching strategies in higher education to improve student performance and decrease drop-out, especially in mathematics, engineering or science courses (Freeman et al., 2014), including population genetics and evolution (Soderberg & Price, 2003). Practical lessons involving *ad hoc* case studies, interactive visualizations and hands-on laboratory experiments bring abstract concepts to life and are an important part of teaching and learning in higher education scientific-technical disciplines (Dunne & Brooks, 2004). Case studies can help undergraduate and graduate students gain a more practical perspective on what they are learning (Yin, 2009), enabling them to develop critical-thinking, problem-solving skills and a deeper understanding of scientific concepts and processes (Mahdi et al., 2020).

Since the late 1990’s and 2000’s, a large amount of population genetics analysis computer programs were developed by scientists around the world, with the aim of properly exploiting the steadily growing amount of data available due to the development and affordability of molecular marker techniques. Indeed, some of them became very popular, such as GenePop (Raymond and Rousset, 1995), PopGene (Yeh et al., 1999) or Arlequin (Excoffier et al., 2005) to calculate diversity and differentiation parameters; SGS (Degen et al., 2001) or SPAGeDi (Hardy & Vekemans, 2002), for spatial genetic structure at different scales; Cervus (Marshall et al., 1998), TwoGener (Smouse et al., 2001) or Famoz (Gerber et al., 2003) for paternity and parentage analysis. These software tools, originally designed for research purposes, can be used also for a practical approach in population genetics education. Nevertheless, non-user-friendly input requirements often hampered their wide application in the classroom.

More recently, other R-programming language-based population genetics applications were developed, such as PopGeneReport (Adamack & Gruber, 2014) or POPPR (Kamvar et al., 2014); even some of them were adapted to be applied in an educational context, such as LearnPopGen (Revell et al., 2019) or in the collaborative space popgen.nescent (Kamvar et al., 2017). However, using these resources requires some previous programming skills that, although desirable, are not always the case among students. Other ready-to-use applications have been developed implementing many of these analysis in a very accessible way, as the MS Excel Add-in GcenAlEx (Peakall & Smouse, 2012), facilitating therefore the application of practical approaches in population genetics teaching.

Nevertheless, while it has been recognized the importance of theoretical concepts to design, select, perform and interpret case studies (Keen & Packwood, 1995), providing the adequate datasets for practical exercises is always a dull and unappreciated task for the teacher, crucial to reach the learning objectives (Stake, 1995). Therefore, datasets should be carefully designed to illustrate research questions and to help develop or refine theory (Crowe et al., 2011). Nowadays, an overwhelmingly increasing amount of genetic and genomic data derived from open science research projects are available in public databases. Very often, however, these real datasets are not suitable to illustrate a particular theoretical concept, and simulated virtual datasets are required to reach learning objectives. Indeed, in order to evaluate the evolution of one or several populations, the analytical solutions should incorporate a multi-generational approach to monitor the population histories through “snapshots”, as if they were independent populations. Simulation of virtual populations arises as an appropriate approach for this task. This is the case of GENIE (Castillo et al., 2022), focused on genetic drift on populations formed by individuals defined by a single haploid locus, or QGSHINY (Neyhart & Watkins, 2020), aimed to illustrate concepts of quantitative genetics under simple demographic scenarios. Other practical approaches that allow monitoring of populations have been applied to teaching population genetics (e.g. Revell, 2019), these software tools are focused on the changes in time of population genetics parameters, without enabling the possibility of creating specific genotype datasets to answer to different questions.

In this work, we present SimHyb 2, a software to simulate the evolution of hybridizing populations (or species) under user-defined conditions. SimHyb 2 provides a broader scope, and constitutes a useful tool to explore and illustrate the effect of the main evolutionary forces (migration, drift, selection, reproductive success), particularly suitable when dealing with species for which direct experimentation is not feasible (such as long-living organisms as trees or certain animal species, for instance). Contrarily to other software, such as HybridLab (Nielsen et al., 2006), which produces genotypes only based on allele frequencies, SimHyb 2 works with virtual individuals, allowing them to mate under user-defined conditions. It is the only software, to our knowledge, providing a thoroughly traceable pedigree for each virtual individual. Furthermore, the output of SimHyb 2 can be coupled with other population genetics popular software tools, such as GenAlEx.

SimHyb 2 is particularly appropriate for practical methodologies, which are increasingly acknowledged in teaching, such as Project Based Learning (PBL) (Kokotsaki et al., 2016) or Directed Studies (DS) (Moore et al., 2018). Students can produce their own genotype datasets with SimHyb 2, and use them directly in further analyses in the project. Due to the stochastic procedures included in different phases of the simulation process, outputs will never be identical (in terms of genotypes, pedigrees, survival…). This uniqueness of the case studies is also a desirable feature from the teacher’s point of view, to enable further group discussion or cooperative learning methodologies.

### SimHyb 2 functioning

SimHyb 2 is programmed in Java 16, and runs in any computer with an OS that allows for Java (https://www.java.com/), or OpenJDK (http://openjdk.java.net/): Linux/Unix, Microsoft Windows, Mac OS X, and other platforms, including cloud virtual desktops that operate in academic institutions to facilitate e-learning (Moser et al., 2014). The application provides a user-friendly front-end to facilitate its using by the students. The software, installation instructions, user manual and input examples are available at https://github.com/GGFHF/SimHyb2

#### Overview

SimHyb 2 simulates the evolution of a population of constant size with up to two different diploid genetic groups (we will refer to them as “species”, for simplicity), which may hybridize, depending on the conditions defined by the user. The population consists of a fixed number of individuals, each of them identified by a vector including pedigree, a so-called “specific coefficient” (representing the expected contribution of each species to the individual genome), the complete genotype for a user-defined number of loci and other information. Initially, SimHyb 2 was intended for long-living plants, such as trees (so it considers features such as the presence of chloroplast, overlapping generations or the possibility of self-incompatibility processes), but it can be also applied for other organisms, taking into account certain considerations discussed below.

In each cycle, in order to produce offspring, pairs of individuals are selected randomly for mating. Individuals are considered hermaphrodite, so they can act both as mother and as father. If the parent pair passes the species-specific fertility and the self-incompatibility filters, a viable offspring individual is produced, and its genotype is defined, drafting alleles from the parents. A fitness value is also assigned to the individual, based on its specific composition and on the particular parental values. According to user specifications, immigrant individuals can also incorporate to the population at this stage.

Then, individuals from previous cycles are aged, reducing their fitness values. After ageing, individuals with the lowest fitness are removed, so that the population remains constant. Finally, before initializing the next cycle, fitness values of survivors are standardized between 0 and 1. Overview of SimHyb 2 functioning is depicted in Figure 1.

**Figure 1.**
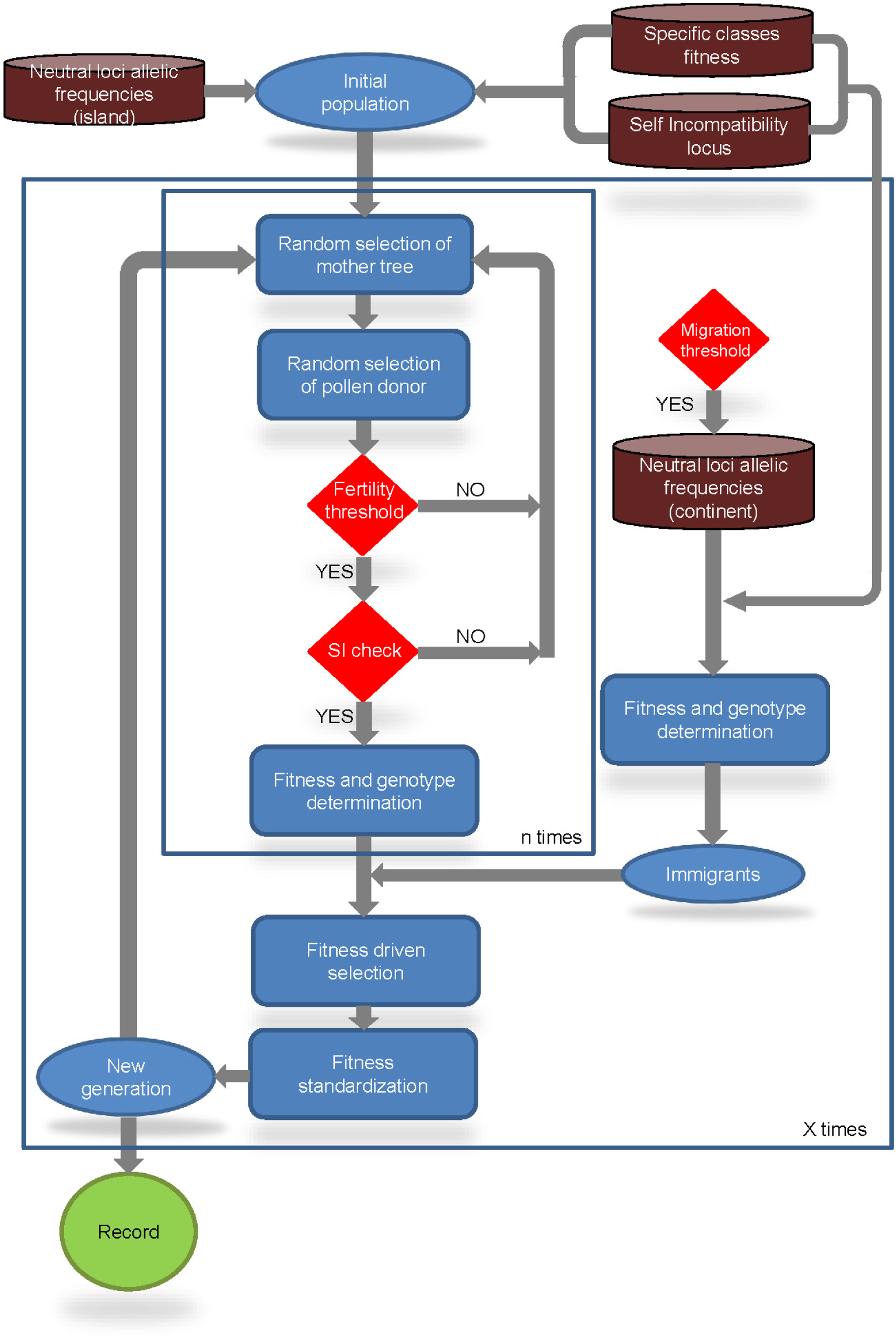
Overview of the workflow of SimHyb2.

#### Virtual individuals

SimHyb 2 operates with virtual individuals defined by the following attributes: (1) a nuclear genotype consisting of a variable number of diploid neutral loci and a diploid self-incompatibility locus; (2) two organellar genotypes (chloroplast and mitochondria), each one considered as a single haplotypic locus, associated to the species; (3) a specific coefficient (representing the expected contribution of each species to the individual genome); and (4) the ID of the parents of each individual.

#### Input files

Before starting the simulation, an interface window appears where the user must define the simulation parameters and input files (Figure 2). The Instructions Manual provided with the software includes examples of these input files (https://github.com/GGFHF/SimHyb2).

**Figure 2.**
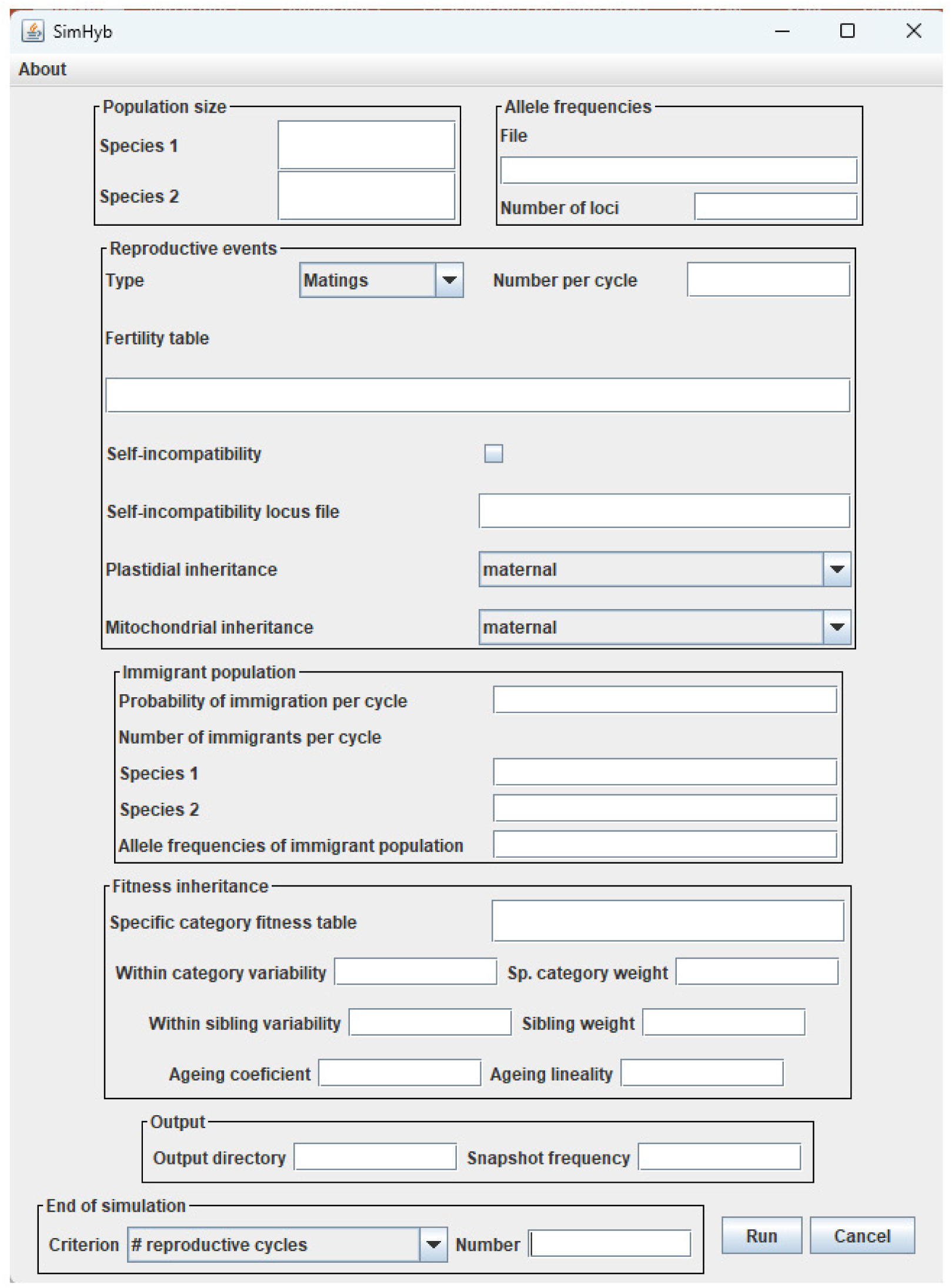
SimHyb 2 interface.

##### Self incompatibility locus

A *.csv* file including the alleles (first column), their frequencies in species 1 (second column) and their frequencies in species 2 (third column). If one allele is absent in one of the species, a *“0”* value must be recorded in the corresponding position.

##### Neutral loci

SimHyb 2 admits up to 1,000 neutral diploid co-dominant loci. Allele frequencies are also provided as a *.csv* input file. The file consists on a rectangular matrix. The left half of the matrix corresponds to the species 1 and the right half to species 2. All the loci must be present in both species (even as monomorphic ones), and reported in the same order. For each species, every locus is represented by two columns: the first one includes the alleles and the second one their frequencies in that species. The same alleles must be included for both species, recording a 0-frequency value where needed. As not all the loci necessarily have the same number of alleles, *“0”* must be included where needed in order to get a rectangular matrix.

##### Specific category and Fitness

An arbitrary, numeric *“specific coefficient”* is assigned to each of the two species included in the initial population. The user can define as many intermediate specific classes as desired, with correspondingly intermediate specific coefficient values. An average fitness value, between 0 and 1, is also assigned to each specific category. These data are provided as a .txt file. Each row corresponds to a specific category, with elements separated by “*;”*. It includes a correlative cardinal, the name of the category (f. i. *species 1, hybrid 1, hybrid 2…*), the lower and upper limits of the specific coefficient for the category (integer numbers are recommended), and the category fitness value, between 0 and 1.

##### Fertility matrix

Fertility among the defined specific categories, which takes into account the pollination direction, is also defined as a *.txt* input matrix, with the specific categories acting as male parents as columns and as female parents in rows.

#### Initial population

The user can define the initial population size for each species, considering only adult individuals. The size of the global population will remain constant throughout the simulation. Genotypes of the individuals of the initial population are generated drafting alleles according to their frequencies in each species, from the input files described above. A chloroplast and a mitochondrion are assigned to each species, and recorded for each individual. *“Null”* values are recorded as the parents of the initial population individuals. Individual fitness values are assigned randomly from the continuous interval *(f_sp_-ε_sp_, f_sp_+ε_sp_)*, being *f_sp_* the average value for the specific category and *ε_sp_* the within category variability defined by the user.

#### Reproductive events

User can define the type (matings or offspring) and number of reproductive events to perform in each cycle. Up to six steps take place to obtain the next generation of adult (reproductive) individuals.

##### Fertility and self-incompatibility filters

For each event (mating or offspring), a random individual is selected as female parent and a second one as male parent. The probability of obtaining a viable cross is firstly determined by the fertility coefficients defined in the input file fertility.txt, according to the specific classes of the female parent and the male parent. After that, effective mating is limited by self-incompatibility, if this option is selected. SimHyb 2 considers gametophytic self-incompatibility driven by a single locus. Doing so, one of the alleles at this locus from the male parent is randomly selected and compared with both alleles of the female parent. The offspring is produced only if no coincidence is registered. The species coefficient of the new individual will be calculated as the average of the parents’ values, so it can be used as a precise estimation of the contribution of species to the genome of the individual.

##### Offspring genotype

For the self-incompatibility locus, the new individual will carry the paternal allele previously selected and one from the mother, chosen at random. It will also inherit at each neutral diploid locus one allele from each parent, randomly selected. Organelles (chloroplast and mitochondria) are inherited from the parents according to the user specifications.

##### Fitness inheritance

A fitness coefficient will be assigned to the new individual as a weighted average of a specific and a family component, according to the following formulae:

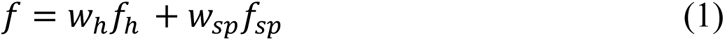

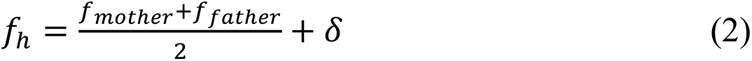

where *f_sp_* is the fitness coefficient corresponding to the specific category of the new individual, *f_h_* is the fitness inherited from the parents; *δ* is drafted from the interval (-*εh, εh*), being *εh* the *“within sibling variability”* parameter, in order to include variability among full-sibs; *w_h_* and *w_sp_* are the corresponding weights, established by the user, so that *w_h_* +*w_sp_* = 1.

##### Immigration

At this point, immigrant individuals can be incorporated into the population, according to a user-defined probability (between 0 and 1). The user also defines the number of individuals in the immigrant pool (they can be different number for each species). The current version of SimHyb 2 only considers the *“continent-island”* migration model. Thus, the evolving population would be the *“island”*, which can incorporate immigrants from a *“continent”* whose allelic frequencies do not vary during the simulation. Immigrants are created in the same way that the initial population, but a *.csv* input file with the allelic frequencies for the *continent* population, with the same loci and alleles (it can be the same file as for the *island*).

##### Ageing

Once the defined number of reproductive events (matings or offspring) has been achieved, individuals from previous cycles are aged, reducing their fitness values, according to this formula:

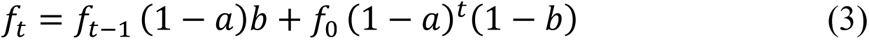

where *f_t_* is the fitness coefficient *t* cycles after birth, *f_0_* is the initial fitness of the individual, at the time of birth, *a* is the *“ageing coefficient”*, and *b* is the *“linearity coefficient”*, both of them defined by the user, between 0 and 1.

##### Fitness-driven selection and standardization

After ageing, in order to keep the population size constant, individuals with the lowest fitness are removed, and fitness values of survivors are standardized between 0 and 1. Then, a new reproductive cycle initializes.

#### End of simulation and output files

Three alternative criteria can be followed to finish the simulation: (1) a user-defined number or cycles; (2) the presence of individuals of a user-defined number of specific categories in the simulation; and (3) fixation of a chloroplast type in the population. This latter option allows the user to explore the demographic and/or reproductive conditions in which one species’s organelle can be captured by the other species, and even replace completely the original one (as proposed, for instance, for the *Quercus suber* and *Q. ilex* in Eastern Spain and Southern France; Jiménez et al., 2004). Additionally, the user can define a maximum number of cycles as security limit.

SimHyb 2 provides two output files. The first one includes all the individuals in the population every X cycles, as a sort of “snapshot album” of the evolving population. The user can define the frequency of these “snapshots” and the output can be consulted while the simulation is still running. The second output file is available only when the simulation is finished, and includes all the individuals of the population in each cycle.

Output files are provided as *.csv* files. The first row registers the headings of the first 10 columns, which include different information of each individual (see below). The second row includes the name of the nuclear, diploid loci, starting with the SI locus and followed by the neutral loci. Third and successive rows include the virtual individuals. The first 10 positions include the following information: (1) Individual ID (integer); (2) Specific coefficient (numeric character); (3) Father individual ID; (4) Mother individual ID; (5) Chloroplast (A or B); (6) Mitochondria (A or B); (7) Generation (integer; cycle of the simulation); (8) Birth Generation (integer; cycle in which the individual is added to the population); (9) Death Generation (integer; cycle in which the individual is removed from population; *“-1”* for survivors); and (10) individual fitness value in that generation (number, between 0 and 1). The next positions register the diploid genotype of the individual, starting with the self-incompatibility locus (alphanumeric characters, two alleles) and neutral loci (alphanumeric characters, two alleles per locus).

Therefore, this output file is easily convertible into the input file of different and widely used population genetics software tools. For instance, just modifying the heading rows and removing some of the first columns would be needed to get an input file for GenAlEx (Peakall & Smouse, 2012) or NewHybrids (Anderson & Thompson, 2002). In a similar way, removing the first row is enough to get an input file for Structure (Pritchard et al., 2000), where columns 3-10 or 3-12 can be labelled as “extra columns”. Of course, non-desired rows (according to the analysis to be performed), such as repeated individuals, present in the population for several cycles (where the only variation would be registered in the relative fitness value), must also be removed accordingly.

### Application of SimHyb 2. Case studies

One of the main applications of SimHyb 2 is to illustrate the effects of evolutionary forces on the genetic pool and allelic frequencies of a population, and the generation of appropriate datasets for academic exercises and further analysis with other popular software tools such as GenAlEx or Structure. Other commonly used software, such as HybridLab (Nielsen et al., 2006), only allows the production of virtual F1 or F2 hybrid individuals. It drafts alleles according to their frequencies in the parental populations (species) to complete the genotype of hybrid individuals, and requires the calculation of frequencies in the hybrid output to be included as an input in a following cycle if further introgression levels are desired. On the contrary, SimHyb 2 uses the original allele frequencies only in the construction of the first generation of the population, and later on, those individuals reproduce among them, according to user-defined rules. In doing so, SimHyb 2 provides traceable pedigrees of individuals with known, different introgression levels. Therefore, SimHyb 2 outputs provide the genotypes and information to prepare datasets for exercises on parentage analysis, on spatial genetic structure, or for Conservation Genetics practical cases. SimHyb 2 can also be used for research purposes, as did the previous version, to assess hybridization and introgression processes (Soto et al., 2018; López de Heredia et al., 2018, 2020) or to check suitability of markers for different purposes (Cosín-Roldán et al., 2023).

SimHyb 2 is intended for long-living plants, with over-lapping generations. Nevertheless, user can set a large ageing coefficient (f.i., *a = 1*) in order to get “annual” (or, at least, non-overlapping) generations. Notwithstanding, application for animal species is more difficult, not only due to the inclusion of chloroplast (which could be simply overlooked), but mainly because individuals are considered hermaphrodite (not appropriate for most animals).

Four examples of application of SimHyb 2 are shown below. Simulation conditions are detailed in Supplementary Information. The ongoing use of SimHyb 2 in under- and postgraduate teaching is also reported.

#### Case study 1: Selective advantage

Here, SimHyb 2 has been used to simulate the evolution of a mixed population of two non-hybridizing species, with species 2 showing higher fitness than species 1. As expected, species 2 outcompetes species 1 during the simulation, leading to the extinction of species 1 in cycle 16. This effect can also be appreciated in the evolution of global allelic frequencies (Figure 3).

**Figure 3.**
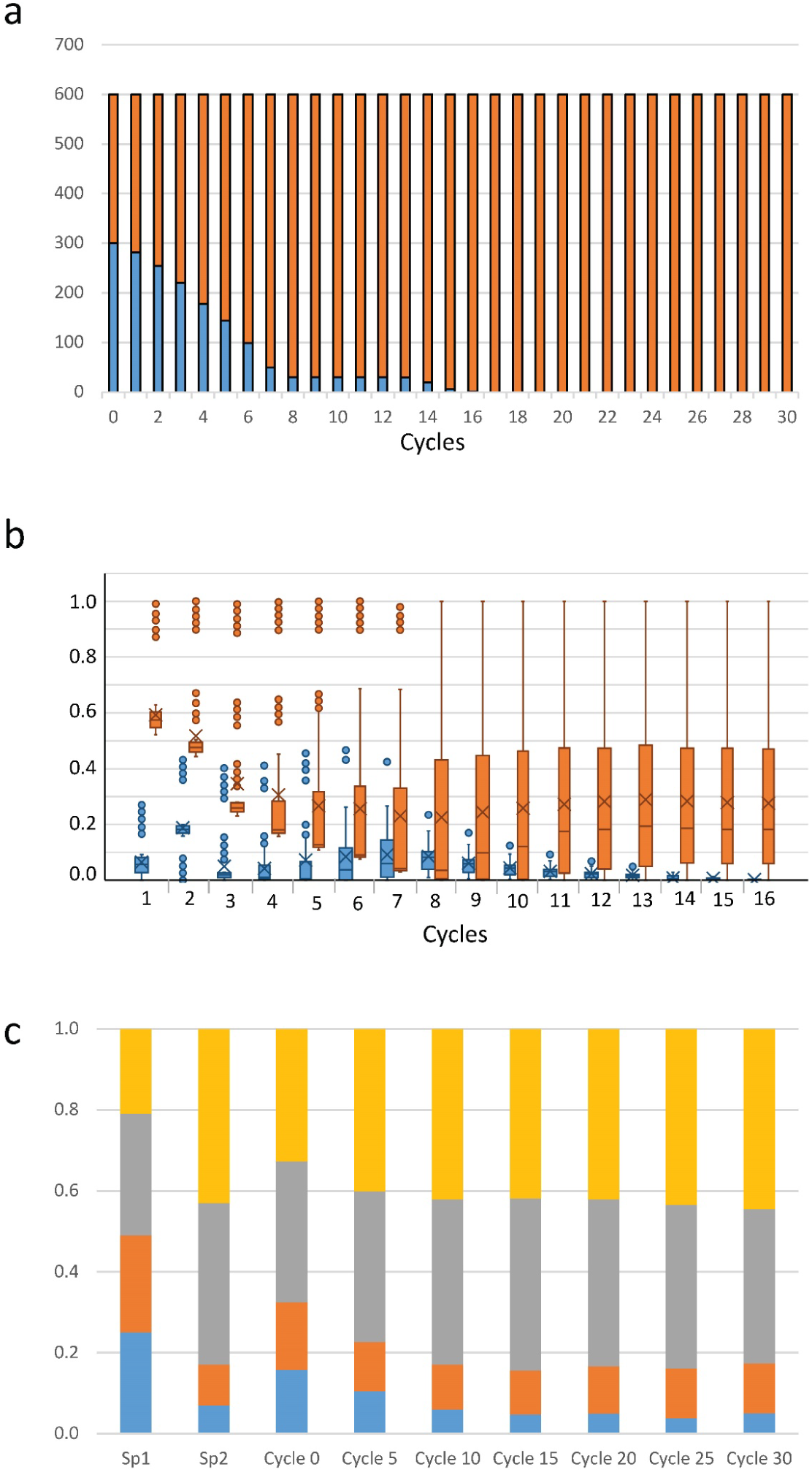
a) Evolution of the number of individuals from species 1 (blue bars) and species 2 (orange bars) in the mixed population. b) Evolution of the relative fitness values per species (sp1 in blue, sp2 in orange) in the first 16 cycles, standardized between 0 and 1 in each cycle for the total mixed population. In addition to the lower values initially assigned to this species, ageing of surviving individuals leads to the final extinction of species 1 in cycle 17. c) Evolution of the global allelic frequencies (for the total mixed population) for two loci throughout 30 reproductive cycles. Initial allele frequencies for each species are also represented.

#### Case study 2: Migration

In this example we considered again the evolution of a mixed population of two non-hybridizing species (*island*), but assimilating migrants from a reference population (*continent*). Initially, species 1 allele frequencies are quite different in the *island* and the *continent* for loci 1-4, and similar for loci 5-8, while species 2 frequencies are identical in both locations.

Figure 4 shows the evolution of allelic frequencies for species 1 throughout 30 reproductive cycles for three loci, as illustration. Variation for species 2 is almost unappreciable and due basically to genetic drift. On the contrary, species 1 shows important changes, with allele frequencies converging to those of the continent. This effect is also appreciable in the differentiation parameter “pairwise F_ST_”, calculated per species and cycle (Table 1). While pairwise-F_ST_ between the cycles 0 and 30^th^ gets to nearly 10% for species 1, it barely gets to 1% for species 2. F_ST_ between both species increases form nearly 10% at the beginning of simulation up to 16% at the end.

**Figure 4.**
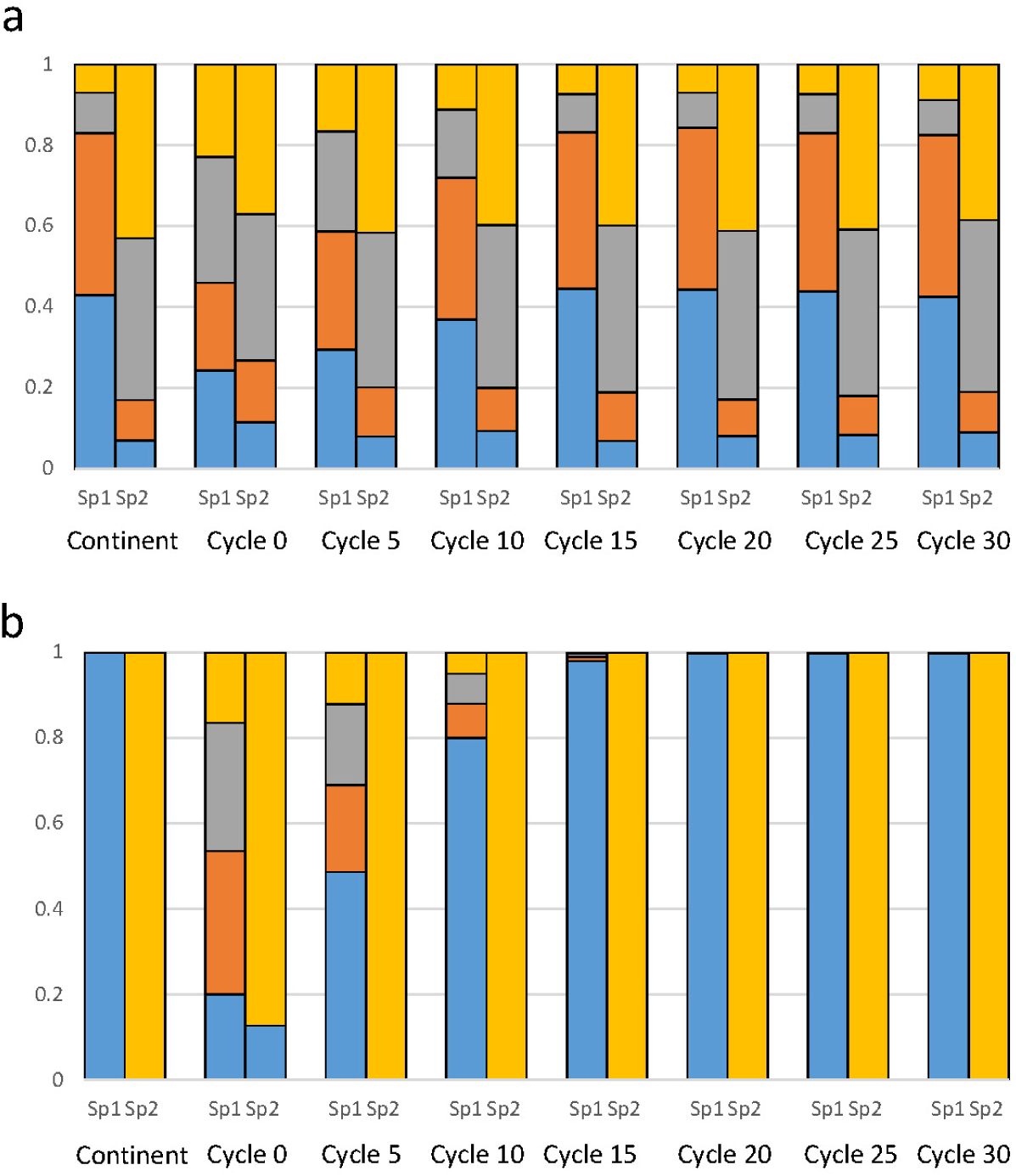
Evolution of the allelic frequencies for two loci (a, b) throughout 30 reproductive cycles both for species 1 and 2. Allele frequencies in the continent are also represented.

**Table 1.** Pairwise-F_ST_ per species and cycle throughout the simulation. While immigration has virtually no effects for species 2, a much larger differentiation can be appreciated in species 1. Between-species differentiation also increases throughout simulation (lower left corner).

| Species | Cycle | Sp 1 |  |  |  | Sp 2 |  |  |  |
| --- | --- | --- | --- | --- | --- | --- | --- | --- | --- |
|  |  | 0 | 10 | 20 | 30 | 0 | 10 | 20 | 30 |
| Sp 1 | 0 | 0.000 |  |  |  |  |  |  |  |
|  | 10 | 0.058 | 0.00 |  |  |  |  |  |  |
|  | 20 | 0.098 | 0.012 | 0.00 |  |  |  |  |  |
|  | 30 | 0.099 | 0.012 | 0.00 | 0.00 |  |  |  |  |
| Sp 2 | 0 | 0.098 |  |  |  | 0.00 |  |  |  |
|  | 10 |  | 0.120 |  |  | 0.009 | 0.00 |  |  |
|  | 20 |  |  | 0.165 |  | 0.010 | 0.001 | 0.00 |  |
|  | 30 |  |  |  | 0.158 | 0.009 | 0.001 | 0.001 | 0.00 |

#### Case study 3: Hybridization / Suitability of markers for introgression analysis

The user can also apply SimHyb 2 to produce hybrid populations, where the specific coefficient of each individual provides the expected contribution of the initial groups/species to its genome. These SimHyb 2 outputs can be easily transformed into Structure (Pritchard et al., 2000) inputs, by simply modifying the first two rows accordingly. Structure is a popular software for genetic classification of individuals to inferred or provided groups or clusters, applying a Bayesian approach, and based on the individual genotypes. Thus, the user can compare the estimations obtained by Structure with the *specific coefficient* provided by SimHyb 2, in order to evaluate the capability of the loci included in the genotypes to detect and infer the individual introgression level.

As an illustration, in this case study we have simulated the evolution of mixed populations, favoring hybridization, for 5 generations, allowing the appearance of up to 18 hybrid classes, in addition to the parental species (Table 2). Individual genotypes included eight independent, neutral loci, with four alleles each.

**Table 2.** Specific categories considered, expected contribution of species 1 to individual genomes and cycle of appearance in the SimHyb 2 simulation. Composition of virtual populations analyzed with Structure.

| **Specific category** | **Specific coefficient (SIMHYB 2)** | **Cycle of appearance (SIMHYB 2)** | **Number of individuals (STRUCTURE)** | **Percentage (STRUCTURE)** |
| --- | --- | --- | --- | --- |
| Species 1 | 1.00 | 1 | 3,100 | 44.50 |
| Hybrid 1 | 0.94 | 5 | 7 | 0.10 |
| Hybrid 2 | 0.88 | 4 | 30 | 0.43 |
| Hybrid 3 | 0.81 | 5 | 30 | 0.43 |
| Hybrid 4 | 0.751 | 5 | 5 | 0.07 |
| Hybrid 5 (backcross) | 0.75 | 3 | 100 | 1.44 |
| Hybrid 6 | 0.69 | 5 | 20 | 0.29 |
| Hybrid 7 | 0.63 | 4 | 50 | 0.72 |
| Hybrid 8 | 0.56 | 5 | 15 | 0.22 |
| Hybrid 9 | 0.501 | 5 | 3 | 0.04 |
| Hybrid 10 (F1) | 0.50 | 2 | 250 | 3.59 |
| Hybrid 11 | 0.44 | 5 | 15 | 0.22 |
| Hybrid 12 | 0.38 | 4 | 50 | 0.72 |
| Hybrid 13 | 0.32 | 5 | 20 | 0.29 |
| Hybrid 14 | 0.251 | 5 | 5 | 0.07 |
| Hybrid 15 (backcross) | 0.25 | 3 | 100 | 1.44 |
| Hybrid 16 | 0.19 | 5 | 30 | 0.43 |
| Hybrid 17 | 0.12 | 4 | 30 | 0.43 |
| Hybrid 18 | 0.06 | 5 | 7 | 0.10 |
| Species 2 | 0.00 | 1 | 3,100 | 44.50 |

Then, virtual populations including individuals with different levels of introgression (Table 1) were analyzed with Structure, estimating the contribution of each parental species to the genome of each individual, considering subsets of four loci. Figure 5 illustrates the comparison of these estimations with the actual expected contributions, provided by SimHyb 2, for each subset. Average values of Structure estimations for each specific category using complete genotypes are fairly close to the expected ones, although individual deviations are large in many cases. Accuracy is provided mostly by the first subset of markers, loci 1-4, while loci 5-8 are completely inappropriate for these purposes.

**Figure 5.**
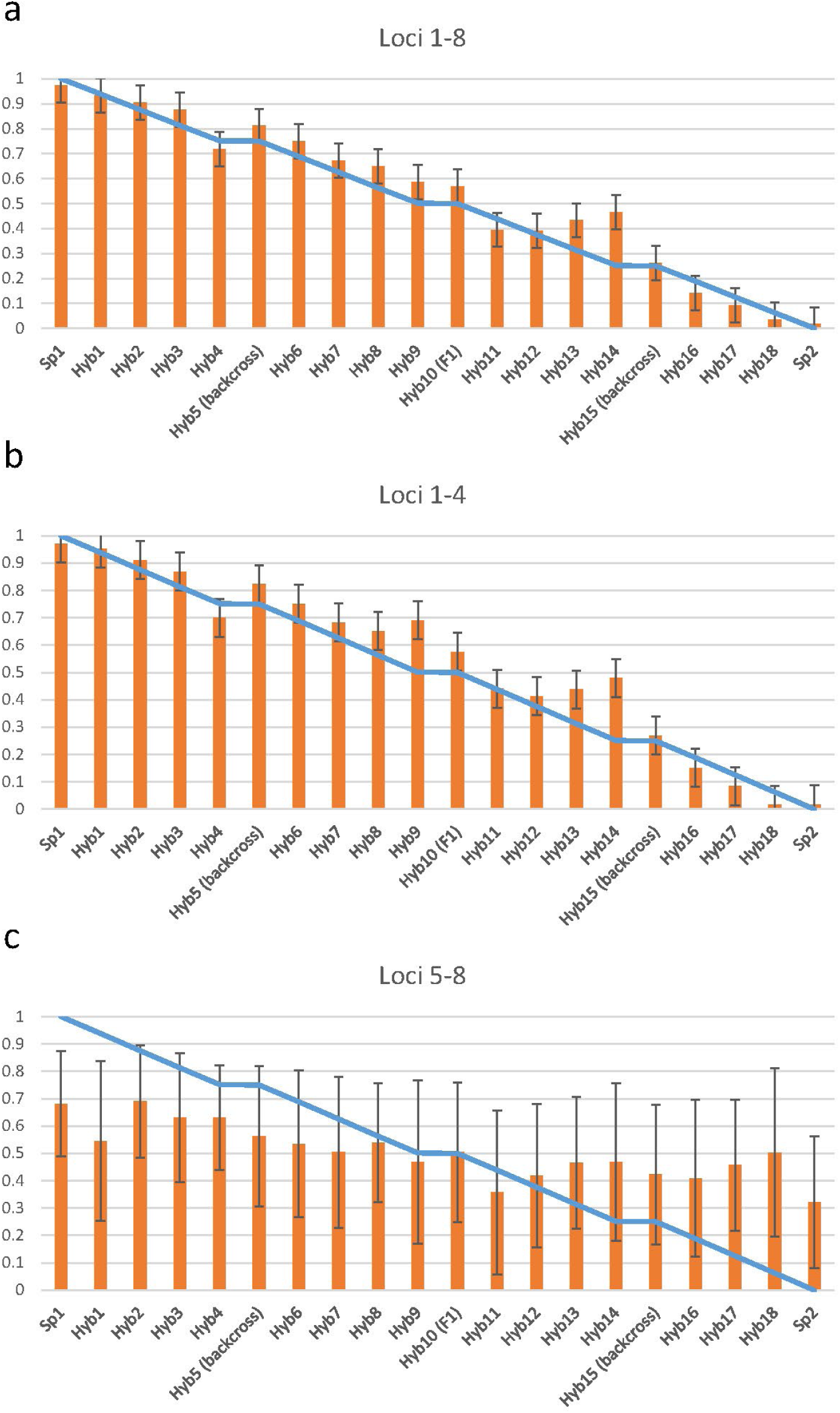
Comparison of expected contribution of species 1 to individual genomes provided by SimHyb 2 simulations (blue line), according to specific categories, and estimations obtained with Structure (orange bars) using a) complete genotypes with eight loci, b) loci 1-4 and c) loci 5-8. Standard errors are shown.

#### Case study 4: One invasive species and directional introgression

In this example, the original population includes individuals from a single species, and individuals from a second species are incorporated from the *continent*. These immigrants display the same fitness as the native species, but they show a higher reproductive success (higher intraspecific fertility value). In addition, hybrids show higher fitness values and backcrossing with the immigrant population are favored, increasing therefore the chances of propagation for immigrant alleles. Population composition and average specific coefficient vary accordingly during the simulation (Figure 6).

**Figure 6.**
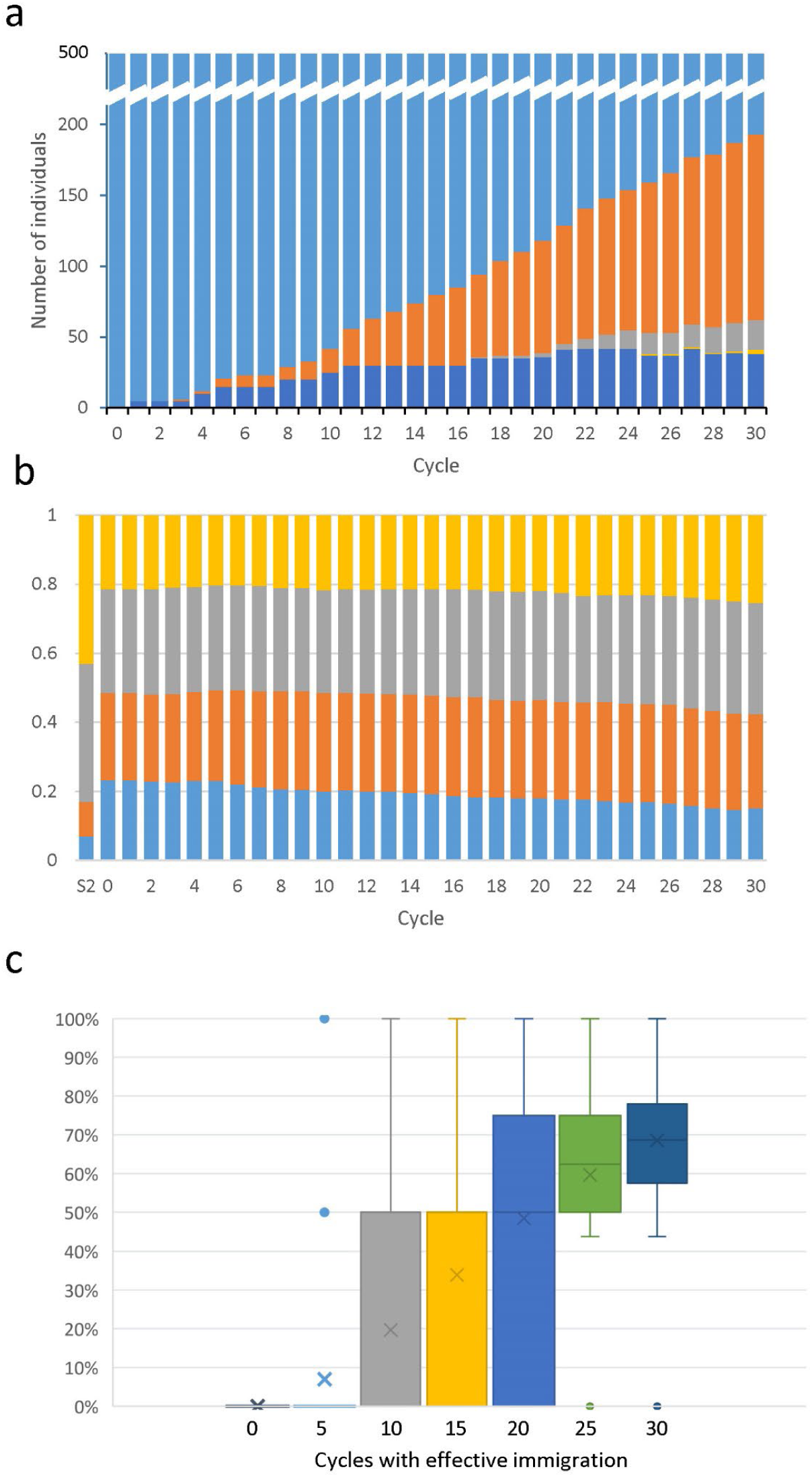
Immigration of an invasive species. a) Evolution of the number of individuals according to specific categories. Species 1 (native, light blue), species 2 (invasive, dark blue), hybrids with different contribution of species 2 to the genome (40-60%, orange; 60-80%, grey; 80-100%, yellow). b) Evolution of the allelic frequencies in the global population throughout 30 cycles for one locus Allelic frequencies for the invasive species (S2) is also represented. c) Box and whisker plot of the evolution of the average contribution of species 2 to the individual genome throughout 30 cycles of accumulated effective immigration.

#### Combined use of SimHyb 2, GenAlEx and Structure in academic courses and results achieved

In addition to the evolutive cases described above, we have applied SimHyb 2 combined with GenAlEx and Structure for project-based learning in the course “Forest Genetic Resources Conservation and Breeding” (Master in Forest Engineering; Universidad Politécnica de Madrid) (autumn semester) during the courses 2021-2022, 2022-2023 and 2023-2024. Most of the mark of the course (70%) is achieved through practical work, in which students use the software extensively. First, students run simulations under different scenarios of fitness, reproductive success, hybridization, migration, genetic drift. SimHyb 2 outputs are then analyzed with other software (as GenAlEx) and modification of allelic frequencies, diversity and differentiation parameters, etc. throughout the evolution of simulated populations are discussed, illustrating this way the effect of evolutionary forces. As well, students estimate introgression levels with Structure and compare them with the actual values provided by SimHyb 2; this practice allows students to take in the relevance of selecting the appropriate markers for different purposes. After this sort of training period, a larger project is undertaken. SimHyb 2 is used to generate virtual individuals with known introgression levels between two “species” or genetic pools. Also, long SimHyb simulations are run using the same initial conditions and frequencies, in order to generate virtual populations derived from the same ancestral one. These individuals are combined to produce *ad hoc* virtual populations, which are located throughout the distribution range of a putative species or pair of species. These genotypes are provided to the students, as corresponding to a sampling, together with information regarding land ownership, protection status, economic value, land use, threads, etc. The students, in groups, must analyze these SimHyb-produced populations with GenAlEx, obtaining their basic genetic parameters, distances, etc. In addition, they estimate the number of genetic pools in the distribution area, using Structure, and the population and individual introgression levels. Students use all this information to propose a plan for the conservation of the species genetic resources, combining *ex situ* and *in situ* activities. The plans are presented and discussed in the classroom. This project is aimed to consolidate the knowledge acquired on population genetics and evolution, the interpretation and use of genetic parameters and estimators and their application to realistic challenges in Conservation of Genetic Resources. Finally, in a separate activity, phenotypic data are assigned to virtual individuals with known pedigrees, obtained with SimHyb 2, thus simulating data from field trials; students must then determine significant factors influencing the quantitative variable (blocks, species, families, environmental variables, interactions…), and estimate heritability, feasibility of breeding programs, etc. The results have been very satisfactory, with success rates (passes/evaluated students) of 78.9% in 2021-2022, 87.5% in 2022-2023 and 89.5% in 2023-2024.

At the end of the autumn semester 2023-2024, we have performed an anonymous survey among the 21 students of the course, including 10 questions focused on the usability and usefulness of the software, and the effect on the acquisition and consolidation of their understanding of population genetics and evolution (Table 3). Feedback was very favorable about the performance of SimHyb 2 and especially about the combination with the other downstream applications (GenAlEx and Structure) to achieve the academic goals (Figure 7). Although students identified the handling of input and output files as the major drawbacks of SimHyb 2, students have declared their overall satisfaction with this tool. 75% of students found very effective the presentation and discussion of results for the acquisition of knowledge and skills. Only 15% of students would have preferred a more theoretical approach to the study of genetic evolution of populations, while results were absolutely balanced regarding Conservation Genetics: one third stated that they would have preferred a more theoretical approach, another third disagreed with this option, and the final third were indifferent to this possibility. In any case, 85% of the students felt they learnt more with the group work than from individual tasks and 57% strongly agreed with this statement.

**Figure 7.**
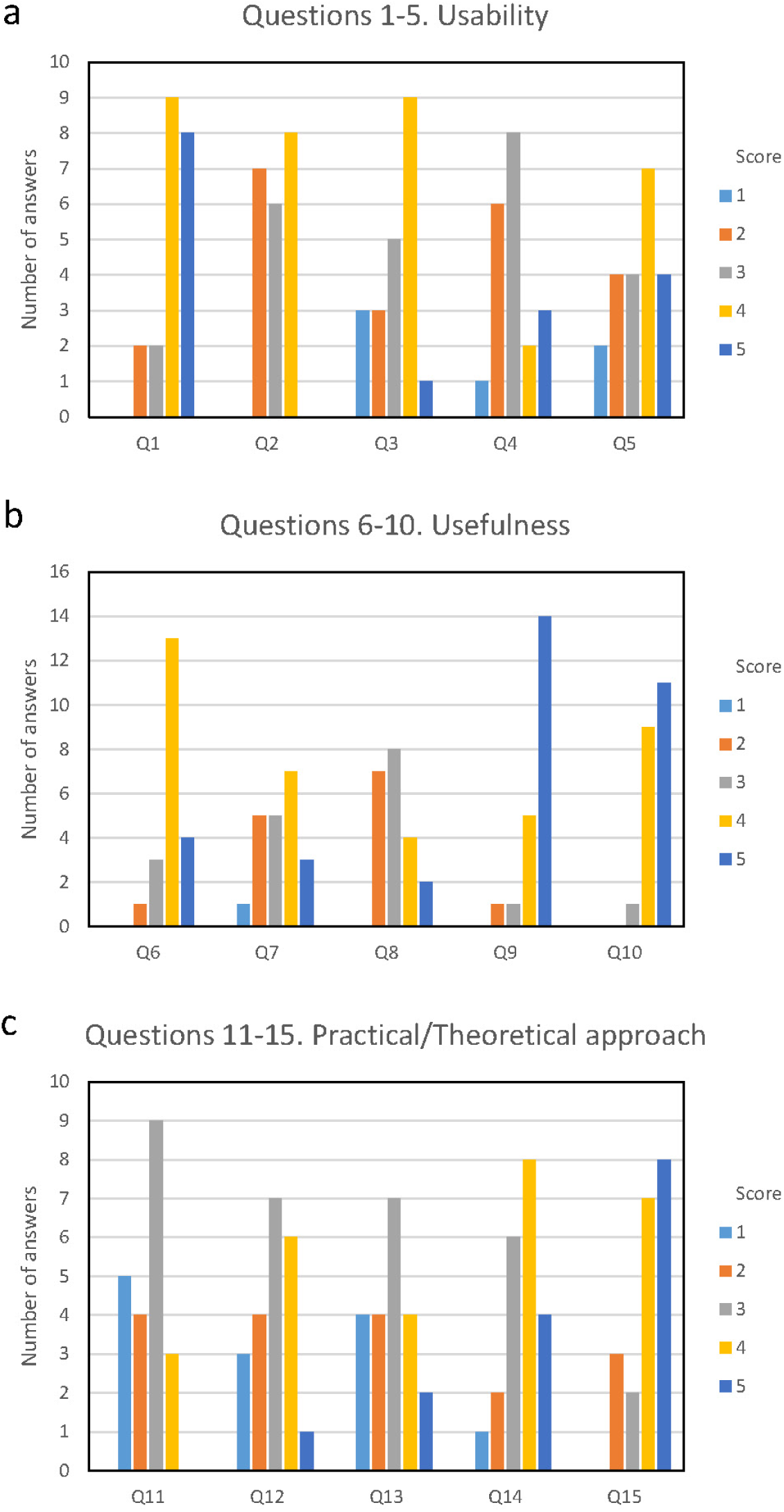
Results of the anonymous survey performed by master students in 2023-2024, including questions related to a) usability of SimHyb 2, compared to other downstream applications (GenAlEx and Structure) (questions Q1 to Q5), b) usefulness for the accomplishment of academic goals (questions Q6 to Q10) and c) educational approach (questions Q11 to Q15).

**Table 3.** Questions included on the anonymous survey regarding the usability (questions Q1 to Q5) and usefulness (questions Q6 to Q10) of SimHyb 2. Students scored from 1 to 5 each aspect.

|  |  |
| --- | --- |
| With 1 being the worst score and 5 the best, rate the following aspects: |  |
| Q1 | Usability of GENALEX |
| Q2 | Usability of STRUCTURE |
| Q3 | Usability of SIMHYB2 |
| Q4 | STRUCTURE interface |
| Q5 | SIMHYB2 interface |
| Rate from 1 to 5 your agreement or disagreement with the following statements, with 1 being "I strongly disagree" and 5 being "I completely agree": |  |
| Q6 | SIMHYB2 correctly simulates different evolutionary scenarios. |
| Q7 | SIMHYB2'S preparation and handling of input files is comfortable and simple |
| Q8 | Modification of SIMHYB2 output files for use with other applications such as GENALEX and STRUCTURE is convenient and straightforward |
| Q9 | The combination of GENALEX, SIMHYB2 and STRUCTURE is useful to illustrate the effects of the different evolutionary forces |
| Q10 | Overall, my knowledge of the genetic evolution of populations has increased. |

These results are roughly consistent with other surveys regarding Genetics teaching and learning, as the one reported by Giménez *et al*. (2021) for a practical approach to strengthen the learning of Mendelian genetics using an *ad hoc* obtained population of barley. Nevertheless, the students in the present survey were more favorable to the advantages of practical approaches than those form Giménez *et al*. (2021). Maybe a more mature attitude can account for this feature: Giménez et al. reported a survey among second course, undergraduate students, 19-20 years old, while the students in the present work were in the second course of a master’s degree, at least 24-25 years old. Students in advanced courses, who often combine their studies with paid jobs, tend to show a more proactive attitude, with a better use of the time dedicated to study (Cao, 2012; Yun *et al*., 2020).

This result is also consistent with our experience with SimHyb 2 and undergraduate students. We have used this tool in order to obtain data for the practices of the subject “Forest Genetics” (Degree in Forest Engineering; Universidad Politécnica de Madrid) during the course 2022-23 (spring semester). We use these virtual data, combined with real ones, as input for basic population genetic analysis: allelic richness, diversity, genetic distances among individuals and population, differentiation parameters… We also assign geographical coordinates to virtual individuals of known pedigree so that students can estimate spatial genetic structure at a local scale and infer effective propagation distances, perform parentage analyses, etc. The results have also been positive, with a noticeable increase in the global success rate for the subject, from 43% in the course 2021-22 (using real datasets) to 59% in the course 2022-23 (using virtual data, specifically designed). These results confirm that *ad hoc* practices performed with the appropriate data account for a better comprehension of theoretical concepts, although improvement has not been as remarkable as for graduate students.

Finally, we have also carried out a satisfactory pilot experience in the subject “Evolution” (Degree in Biological Sciences; Universidad de Córdoba) during the course 2022-2023 (spring semester), as well as in the doctorate course “Population Genetics and its application to Conservation of Forest Genetic Resources” (Doctorate Program “Science and Technology of Agroforestry Systems; University of Extremadura) during the course 2023-2024.

## Conclusion

SimHyb 2 is a user-friendly tool suitable for illustrating the effect of evolutionary driving forces in the dynamics and allele frequencies of populations, especially appropriate for project-based-learning approaches or to implement directed studies. SimHyb 2 outputs can be easily coupled with other commonly used population genetics tools, such as GenAlEx or Structure, expanding the possibilities for teaching population genetics. Since it provides traceable pedigrees and actual values of expected introgression for virtual individuals, it can be also used, both in teaching and in research, for parentage analysis or to explore hybridization processes or test suitability of a marker set for these purposes. Students can use SimHyb 2 in their PCs or through academic virtual desktops either in autonomous or guided mode by the teacher, enriching their learning in population genetics. Future versions of the software should implement features as the possibility of sporophytic self-incompatibility or, above all, the consideration of dioecy, which would facilitate the application to animal species, for example.

## Declarations

## Ethical approval

Not applicable.

## Informed consent

Not applicable.

## Research involving human participants and/or animals

Not applicable.

## Consent to participate

Not applicable.

## Consent to publish

Not applicable.

## Author’s contribution

Conceptualization: Álvaro Soto; Methodology: Álvaro Soto; Software: David Rodríguez-Martínez; Formal analysis and investigation: Álvaro Soto, David Rodríguez-Martínez, Unai López de Heredia; Validation: Álvaro Soto, Unai López de Heredia; Writing – original draft preparation: Álvaro Soto; Writing – review and editing: Álvaro Soto, Unai López de Heredia; Visualization: Álvaro Soto; Supervision: Álvaro Soto, Funding Acquisition: Álvaro Soto; Project Administration: Álvaro Soto. All authors have read and agreed to the published version of the manuscript.

## Availability of data and materials

SimHyb 2 software, user manual and input example are freely available at https://github.com/GGFHF/SimHyb2

