## Supplementary Information for "SIMHYB 2: a software tool to explore and illustrate evolutionary forces in Population Genetics teaching and research. Application to Conservation Genetics"

Case Study 1: ≠Fitness w.o. migration, hybridization

**Specific categories**

| | specific coefficient ran[illegible] | | sp fitness |
|---|---|---|---|
| Sp 1 | 0 | 100 | 0.5 |
| Hybrid 1 | 100 | 200 | 1 |
| Hybrid 2 | 200 | 300 | 1 |
| Hybrid 3 | 300 | 400 | 1 |
| Hybrid 4 | 400 | 500 | 1 |
| Hybrid 5 | 500 | 600 | 1 |
| Sp2 | 600 | 700 | 1 |

| **Fertility** | | Pollen donor | | | | | | |
|---|---|---|---|---|---|---|---|---|
| | | Sp 1 | Hybrid 1 | Hybrid 2 | Hybrid 3 | Hybrid 4 | Hybrid 5 | Sp2 |
| Mother tree | Sp 1 | 1 | 0 | 0 | 0 | 0 | 0 | 0 |
| | Hybrid 1 | 0 | 0 | 0 | 0 | 0 | 0 | 0 |
| | Hybrid 2 | 0 | 0 | 0 | 0 | 0 | 0 | 0 |
| | Hybrid 3 | 0 | 0 | 0 | 0 | 0 | 0 | 0 |
| | Hybrid 4 | 0 | 0 | 0 | 0 | 0 | 0 | 0 |
| | Hybrid 5 | 0 | 0 | 0 | 0 | 0 | 0 | 0 |
| | Sp2 | 0 | 0 | 0 | 0 | 0 | 0 | 1 |

**SimHyb.properties**

Sp1Name=species 1
Sp2Name=species 2
populationSizeSp1=300
populationSizeSp2=300
lociNumber=8
alleleFrequenciesFile=(path) .csv file
selfIncompatibilityLocusFile=(path) .csv file
reproductiveEventsType=offspring
reproductiveEventsNumber=50
specificCategoryFile=(path) .txt file
fertilityTableFile=(path) .txt file
selfIncompatibility=true
plastidialInheritance=maternal
mitrochondrialInheritance=maternal
SpCategoryWeight=1
siblingWeight=0
withinCategoryVariability=0.05
withinSiblingVariability=0.05
ageingCoeficient=0.25
ageingLinealityCoeficient=0
immigrationProbability=0
immigrantsNumber1=0
immigrantsNumber2=0
immigrantAlleleFrequenciesFile=(path) .csv file
outputDirectory=(path)
snapshotFrequency=5
endOfSimulationCriterion=reproductiveCyclesNumber
simulationEndLimit=31
securityLimit=150

Case Study 2: Migration w.o. hybridization; = fitness

**Specific categories**

| | specific coefficient rang | | sp fitness |
|---|---|---|---|
| Sp 1 | 0 | 100 | 0.7 |
| Hybrid 1 | 100 | 200 | 1 |
| Hybrid 2 | 200 | 300 | 1 |
| Hybrid 3 | 300 | 400 | 1 |
| Hybrid 4 | 400 | 500 | 1 |
| Hybrid 5 | 500 | 600 | 1 |
| Sp2 | 600 | 700 | 0.7 |

| **Fertility** | | Pollen donor | | | | | | |
|---|---|---|---|---|---|---|---|---|
| | | Sp 1 | Hybrid 1 | Hybrid 2 | Hybrid 3 | Hybrid 4 | Hybrid 5 | Sp2 |
| Mother tree | Sp 1 | 1 | 0 | 0 | 0 | 0 | 0 | 0 |
| | Hybrid 1 | 0 | 0 | 0 | 0 | 0 | 0 | 0 |
| | Hybrid 2 | 0 | 0 | 0 | 0 | 0 | 0 | 0 |
| | Hybrid 3 | 0 | 0 | 0 | 0 | 0 | 0 | 0 |
| | Hybrid 4 | 0 | 0 | 0 | 0 | 0 | 0 | 0 |
| | Hybrid 5 | 0 | 0 | 0 | 0 | 0 | 0 | 0 |
| | Sp2 | 0 | 0 | 0 | 0 | 0 | 0 | 1 |

**SimHyb.properties**

Sp1Name=species 1
Sp2Name=species 2
populationSizeSp1=300
populationSizeSp2=300
lociNumber=8
alleleFrequenciesFile=(path) .csv file
selfIncompatibilityLocusFile=(path) .csv file
reproductiveEventsType=matings
reproductiveEventsNumber=50
specificCategoryFile=(path) .txt file
fertilityTableFile=(path) .txt file
selfIncompatibility=true
plastidialInheritance=maternal
mitochondrialInheritance=maternal
SpCategoryWeight=0.2
siblingWeight=0.8
withinCategoryVariability=0.01
withinSiblingVariability=0.05
ageingCoeficient=0.6
ageingLinealityCoeficient=0
immigrationProbability=0.5
immigrantsNumber1=50
immigrantsNumber2=50
immigrantAlleleFrequenciesFile=(path) .csv file
outputDirectory=(path)
snapshotFrequency=5
endOfSimulationCriterion=reproductiveCyclesNumber
simulationEndLimit=31
securityLimit=100

Case Study 3: Hybridization, w.o. migration, = fitness. Assesing markers suitability for classification

**Specific categories**

| | specific coefficient ran[illegible] | | sp fitness |
|---|---|---|---|
| Sp 1 | 0 | 100 | 0.7 |
| Hybrid 1 | 100 | 200 | 1 |
| Hybrid 2 | 200 | 300 | 1 |
| Hybrid 3 | 300 | 400 | 1 |
| Hybrid 4 | 400 | 500 | 1 |
| Hybrid 5 | 500 | 600 | 1 |
| Sp2 | 600 | 700 | 0.7 |

**Fertility**

| | | Pollen donor | | | | | | |
|---|---|---|---|---|---|---|---|---|
| | | Sp 1 | Hybrid 1 | Hybrid 2 | Hybrid 3 | Hybrid 4 | Hybrid 5 | Sp2 |
| Mother tree | Sp 1 | 0.7 | 1 | 1 | 1 | 0 | 0 | 1 |
| | Hybrid 1 | 1 | 1 | 1 | 1 | 0 | 0 | 0 |
| | Hybrid 2 | 1 | 1 | 1 | 1 | 0 | 0 | 0 |
| | Hybrid 3 | 1 | 1 | 1 | 0 | 1 | 1 | 1 |
| | Hybrid 4 | 0 | 0 | 0 | 1 | 1 | 1 | 1 |
| | Hybrid 5 | 0 | 0 | 0 | 1 | 1 | 1 | 1 |
| | Sp2 | 1 | 0 | 0 | 1 | 1 | 1 | 0.7 |

**SimHyb.properties**

Sp1Name=species 1
Sp2Name=species 2
populationSizeSp1=700
populationSizeSp2=700
lociNumber=8
alleleFrequenciesFile=(path) .csv file
selfIncompatibilityLocusFile=(path) .csv file
reproductiveEventsType=offspring
reproductiveEventsNumber=500
specificCategoryFile=(path) .txt file
fertilityTableFile=(path) .txt file
selfIncompatibility=true
plastidialInheritance=maternal
mitochondrialInheritance=maternal
SpCategoryWeight=0.2
siblingWeight=0.8
withinCategoryVariability=0.05
withinSiblingVariability=0.05
ageingCoeficient=0.25
ageingLinealityCoeficient=0
immigrationProbability=0.5
immigrantsNumber1=0
immigrantsNumber2=0
immigrantAlleleFrequenciesFile=(path) .csv file
outputDirectory=(path)
snapshotFrequency=5
endOfSimulationCriterion=reproductiveCyclesNumber
simulationEndLimit=5
securityLimit=150

Case Study 4: Invasive species; directional introgression; reproductive success

**Specific categories**

| | specific coefficient ran | | sp fitness |
|---|---|---|---|
| Sp 1 | 0 | 100 | 0.7 |
| Hybrid 1 | 100 | 200 | 1 |
| Hybrid 2 | 200 | 300 | 1 |
| Hybrid 3 | 300 | 400 | 1 |
| Hybrid 4 | 400 | 500 | 1 |
| Hybrid 5 | 500 | 600 | 1 |
| Sp2 | 600 | 700 | 0.7 |

**Fertility**

| | | Pollen donor | | | | | | |
|---|---|---|---|---|---|---|---|---|
| | | Sp 1 | Hybrid 1 | Hybrid 2 | Hybrid 3 | Hybrid 4 | Hybrid 5 | Sp2 |
| Mother tree | Sp 1 | 0.5 | 0 | 0 | 0 | 0 | 0 | 1 |
| | Hybrid 1 | 0 | 0 | 0 | 0 | 0 | 0 | 0 |
| | Hybrid 2 | 0 | 0 | 0 | 0 | 0 | 0 | 0 |
| | Hybrid 3 | 0 | 0 | 0 | 0.5 | 0.7 | 0.7 | 1 |
| | Hybrid 4 | 0 | 0 | 0 | 0.7 | 0.7 | 0.7 | 1 |
| | Hybrid 5 | 0 | 0 | 0 | 0.7 | 0.7 | 0.7 | 1 |
| | Sp2 | 1 | 0 | 0 | 1 | 1 | 1 | 1 |

**SimHyb.properties**

Sp1Name=species 1
Sp2Name=species 2
populationSizeSp1=500
populationSizeSp2=0
lociNumber=8
alleleFrequenciesFile=(path) .csv file
selfIncompatibilityLocusFile=(path) .csv file
reproductiveEventsType=matings
reproductiveEventsNumber=50
specificCategoryFile=(path) .txt file
fertilityTableFile=(path) .txt file
selfIncompatibility=true
plastidialInheritance=maternal
mitrochondrialInheritance=maternal
SpCategoryWeight=0.2
siblingWeight=0.8
withinCategoryVariability=0.01
withinSiblingVariability=0.05
ageingCoeficient=0.6
ageingLinealityCoeficient=0
immigrationProbability=0.5
immigrantsNumber1=0
immigrantsNumber2=5
immigrantAlleleFrequenciesFile=(path) .csv file
outputDirectory=(path)
snapshotFrequency=5
endOfSimulationCriterion=reproductiveCyclesNumber
simulationEndLimit=31
securityLimit=100

NEUTRAL DIPLOID LOCI

| | | | | | | | | | | | | | | | | | | | | | | | | | | | | | | | |
|---|---|---|---|---|---|---|---|---|---|---|---|---|---|---|---|---|---|---|---|---|---|---|---|---|---|---|---|---|---|---|---|
| Island Case Studies 1, 2, 4) | | | | | | | | | | | | | | | | | | | | | | | | | | | | | | | |
| 197 | 0.25 | 213 | 0.1 | 252 | 0 | 110 | 0.2 | 202 | 0.1 | 116 | 0.05 | 246 | 0.3 | 180 | 0.15 | 197 | 0.07 | 213 | 0.3 | 252 | 0.3 | 110 | 0 | 202 | 0.08 | 116 | 0.05 | 246 | 0.27 | 180 | 0.17 |
| 199 | 0.24 | 215 | 0.2 | 253 | 0 | 114 | 0.3 | 204 | 0.1 | 118 | 0.2 | 252 | 0.2 | 183 | 0.26 | 199 | 0.1 | 215 | 0.4 | 253 | 0.25 | 114 | 0 | 204 | 0.12 | 118 | 0.2 | 252 | 0.22 | 183 | 0.24 |
| 201 | 0.3 | 217 | 0.3 | 255 | 0 | 118 | 0.35 | 206 | 0.75 | 120 | 0 | 254 | 0.3 | 186 | 0.31 | 201 | 0.4 | 217 | 0.1 | 255 | 0.4 | 118 | 0 | 206 | 0.74 | 120 | 0.05 | 254 | 0.31 | 186 | 0.33 |
| 211 | 0.21 | 219 | 0.4 | 257 | 1 | 120 | 0.15 | 208 | 0.05 | 122 | 0.75 | 256 | 0.2 | 189 | 0.28 | 211 | 0.43 | 219 | 0.2 | 257 | 0.05 | 120 | 1 | 208 | 0.06 | 122 | 0.7 | 256 | 0.2 | 189 | 0.26 |
| Continent (Case Study 2) | | | | | | | | | | | | | | | | | | | | | | | | | | | | | | | |
| 197 | 0.43 | 213 | 0.2 | 252 | 0.05 | 110 | 1 | 202 | 0.09 | 116 | 0 | 246 | 0.32 | 180 | 0.2 | 197 | 0.07 | 213 | 0.3 | 252 | 0.3 | 110 | 0 | 202 | 0.08 | 116 | 0.05 | 246 | 0.27 | 180 | 0.17 |
| 199 | 0.4 | 215 | 0.1 | 253 | 0.4 | 114 | 0 | 204 | 0.11 | 118 | 0.15 | 252 | 0.18 | 183 | 0.2 | 199 | 0.1 | 215 | 0.4 | 253 | 0.25 | 114 | 0 | 204 | 0.12 | 118 | 0.2 | 252 | 0.22 | 183 | 0.24 |
| 201 | 0.1 | 217 | 0.4 | 255 | 0.25 | 118 | 0 | 206 | 0.7 | 120 | 0.05 | 254 | 0.32 | 186 | 0.32 | 201 | 0.4 | 217 | 0.1 | 255 | 0.4 | 118 | 0 | 206 | 0.74 | 120 | 0.05 | 254 | 0.31 | 186 | 0.33 |
| 211 | 0.07 | 219 | 0.3 | 257 | 0.3 | 120 | 0 | 208 | 0.1 | 122 | 0.8 | 256 | 0.18 | 189 | 0.28 | 211 | 0.43 | 219 | 0.2 | 257 | 0.05 | 120 | 1 | 208 | 0.06 | 122 | 0.7 | 256 | 0.2 | 189 | 0.26 |
| Island (Case Study 3) | | | | | | | | | | | | | | | | | | | | | | | | | | | | | | | |
| 11 | 0.25 | 21 | 0.12 | 31 | 0.48 | 41 | 0.3 | 51 | 0.5 | 61 | 0.4 | 71 | 0.1 | 81 | 0.45 | 11 | 0.75 | 21 | 0.8 | 31 | 0.05 | 41 | 0 | 51 | 0.4 | 61 | 0.3 | 71 | 0.2 | 81 | 0.4 |
| 12 | 0.32 | 22 | 0.15 | 32 | 0.06 | 42 | 0.65 | 52 | 0.2 | 62 | 0.32 | 72 | 0.7 | 82 | 0 | 12 | 0 | 22 | 0 | 32 | 0.35 | 42 | 0 | 52 | 0.1 | 62 | 0.2 | 72 | 0.6 | 82 | 0.1 |
| 13 | 0.18 | 23 | 0.33 | 33 | 0.25 | 43 | 0.05 | 53 | 0.1 | 63 | 0.2 | 73 | 0.15 | 83 | 0 | 13 | 0.25 | 23 | 0.05 | 33 | 0.05 | 43 | 0.45 | 53 | 0.3 | 63 | 0.3 | 73 | 0.2 | 83 | 0.05 |
| 14 | 0.25 | 24 | 0.4 | 34 | 0.21 | 44 | 0 | 54 | 0.2 | 64 | 0.08 | 74 | 0.05 | 84 | 0.55 | 14 | 0 | 24 | 0.15 | 34 | 0.55 | 44 | 0.55 | 54 | 0.2 | 64 | 0.2 | 74 | 0 | 84 | 0.45 |
| Continent (Case Study 3) | | | | | | | | | | | | | | | | | | | | | | | | | | | | | | | |
| 11 | 0 | 21 | 0.05 | 31 | 0.15 | 41 | 0.5 | 51 | 0.2 | 61 | 0.1 | 71 | 0.35 | 81 | 0.2 | 11 | 0.75 | 21 | 0.8 | 31 | 0.05 | 41 | 0 | 51 | 0.4 | 61 | 0.3 | 71 | 0.2 | 81 | 0.4 |
| 12 | 0.25 | 22 | 0 | 32 | 0.15 | 42 | 0.45 | 52 | 0.2 | 62 | 0.45 | 72 | 0.5 | 82 | 0.05 | 12 | 0 | 22 | 0 | 32 | 0.35 | 42 | 0 | 52 | 0.1 | 62 | 0.2 | 72 | 0.6 | 82 | 0.1 |
| 13 | 0 | 23 | 0.15 | 33 | 0.45 | 43 | 0.25 | 53 | 0.5 | 63 | 0.3 | 73 | 0 | 83 | 0.05 | 13 | 0.25 | 23 | 0.05 | 33 | 0.05 | 43 | 0.45 | 53 | 0.3 | 63 | 0.3 | 73 | 0.2 | 83 | 0.05 |
| 14 | 0.75 | 24 | 0.8 | 34 | 0.25 | 44 | 0.25 | 54 | 0.1 | 64 | 0.15 | 74 | 0.15 | 84 | 0.7 | 14 | 0 | 24 | 0.15 | 34 | 0.55 | 44 | 0.55 | 54 | 0.2 | 64 | 0.2 | 74 | 0 | 84 | 0.45 |

| SELF-INCOMPATIBILITY LOCUS | | |
|---|---|---|
| #Allele | Sp1 | Sp2 |
| SI1 | 0.15 | 0 |
| SI2 | 0.17 | 0 |
| SI3 | 0.08 | 0 |
| SI4 | 0.01 | 0 |
| SI5 | 0.14 | 0 |
| SI6 | 0.06 | 0 |
| SI7 | 0.03 | 0.09 |
| SI8 | 0.06 | 0.04 |
| SI9 | 0.05 | 0.1 |
| SI10 | 0.02 | 0.13 |
| SI11 | 0.04 | 0.08 |
| SI12 | 0.12 | 0.02 |
| SI13 | 0.07 | 0.06 |
| SI14 | 0 | 0.08 |
| SI15 | 0 | 0.1 |
| SI16 | 0 | 0.05 |
| SI17 | 0 | 0.08 |
| SI18 | 0 | 0.09 |
| SI19 | 0 | 0.04 |
| SI20 | 0 | 0.04 |
